# Mapping the MOB proteins’ proximity network reveals a unique interaction between human MOB3C and the RNase P complex

**DOI:** 10.1101/2023.05.11.540416

**Authors:** Islam E. Elkholi, Jonathan Boulais, Marie-Pier Thibault, Hong-Duc Phan, Amélie Robert, Lien B. Lai, Denis Faubert, Matthew J. Smith, Venkat Gopalan, Jean-François Côté

## Abstract

Distinct functions mediated by members of the monopolar spindle-one-binder (MOB) family of proteins remain elusive beyond the evolutionary conserved and well-established roles of MOB1A and B in regulating the Hippo pathway. Since MOB proteins are adaptors, understanding how they engage in protein-protein interactions and complexes assembly is essential to define the full scope of their biological functions. To address this, we undertook a proximity-dependent biotin identification (BioID) approach to define the interactomes of all seven human MOB proteins in HeLa and HEK293 cell lines. We uncovered > 200 interactions, of which at least 70% are unreported on BioGrid. The generated dataset reliably recalled the *bona fide* interactors of the well-studied MOBs. We further defined the common and differential interactome between different MOBs on a subfamily and an individual level. We discovered a unique interaction between MOB3C and 7 out of 10 protein subunits of the RNase P complex, an endonuclease that catalyzes tRNA 5’ maturation. As a proof-of-principle for the robustness of the generated dataset, we validated the specific interaction of MOB3C with catalytically active RNase P by using affinity purification-mass spectrometry and pre-tRNA cleavage assays of MOB3C pulldowns. In summary, our data provide novel insights into the biology of MOB proteins and reveal the first interactors of MOB3C, components of the RNase P complex, and hence an exciting nexus with RNA biology.

## INTRODUCTION

The highly conserved mammalian MOB family of proteins is comprised of 7 members and is subdivided into 4 subfamilies: MOB1A/B, MOB2, MOB3A/B/C, and MOB4. This family regulates cell cycle/division dynamics, DNA repair, tissue growth and morphogenesis, in addition to cytoskeletal organization (1–3). These notions globally elected this family as potential players in growth-related disorders such as cancer (4–8). However, these functions are revealed only in the context of the best characterized members, namely MOB1A/B as *bona fide* regulators of the Hippo pathway; followed by the relatively unexplored members, MOB2 as a regulator of NDR kinases activity, and MOB4 as a component of the Striatin-interacting Phosphatase and Kinase (STRIPAK) complex (9–11). The MOB3 subfamily is poorly characterized in terms of binding partners and function beyond speculations based on unvalidated screens’ predictions (1). This might be partially attributed to: (i) the lack of mammalian diversity (absence of orthologues for this subfamily members) in the model organisms where the MOBs were initially discovered and characterized phenotypically such as *saccharomyces cerevisiae* (12,13) (has two MOB proteins: Mob1p and Mob2p) and *drosophila melanogaster* (14) (has four MOB proteins: dMob1, dMob2, dMob3, dMob4) or (ii) considering the 3 MOB3 proteins as a single entity given the structural similarity (15).

MOBs are small ∼20 kDa proteins that overall share 17-96% structural similarity between different members and are thought of as scaffolds or adaptor proteins that mediate their functions mainly through engaging with and assembling protein complexes (1). Therefore, different proteomic approaches aiming at revealing protein-protein interactions (PPIs) have been leveraged to reveal the MOB proteins’ interactomes but mainly within the borders of the Hippo and STRIPAK complexes (11,16,17). Despite the advances from these previous studies, including insights into the crosstalk between MOB1 and MOB4 within the Hippo pathway and STRIPAK complex (17,18), a systematic comparison of the interactomes of all seven MOBs in the same cellular context has not been undertaken before. The work described here addresses this gap.

Here, we harnessed the biotinylation-dependent proximity labelling (BioID) approach (19–21), that bypasses different limitations of standard PPI-profiling techniques (22), to explore the global interactome of the MOB proteins in two different cell lines with an intent to focus on the less characterized MOB3s. We reveal the common and unique interactors among the MOB subfamilies and between subfamily members. Unexpectedly, we discovered that MOB3C exists in the vicinity of 7 out of 10 protein subunits of the RNase P complex, an endoribonuclease that catalyzes the cleavage of 5’ leader from precursor tRNA (23,24). Further investigations confirmed that MOB3C, but not MOB1A, interacts with a catalytically active RNase P complex. Our results provide new insights into the MOB proteins interactors and hence shedding light on their functional diversity beyond the view of being kinase activators. Importantly, we uncover a novel potential connection between MOB3C and RNA biology.

## RESULTS

### BioID proximity labeling screens for mapping the global interactome of MOB proteins

We exploited a BioID pipeline to reveal the MOB proteins’ potential interactors on a global scale to assess how they are functionally related as this remains elusive (2,3). BioID depends on fusing a bait, here the 7 MOBs, to an abortive mutant form of the biotin ligase BirA (BirA*) that stimulates the biotinylation of proteins in approximately 10 nm vicinity of the bait (19). As we and others previously demonstrated (19-21,25), this unbiased approach can: (i) capture transient interactors in addition to those that may be missed due to solubility-related caveats (i.e., insoluble cellular structures and/or use of harsh lysis conditions for disrupting interactions at the cell membrane for instance (22)); and (ii) map the spatial landscape of the bait-specific signaling pathways. To this end, we generated tetracycline-inducible HEK293 and HeLa Flp-In T-REx cells expressing either BirA*-FLAG-MOB (for the seven human MOB proteins; Table S1), BirA*-FLAG, or BirA*-FLAG-EGFP, where the latter two serve as negative controls (Figure 1A). The expression of the different baits (MOB proteins or controls) in HEK293 and HeLa cells as well as the overall cellular biotinylation profiles were validated by western blotting (Figure 1B) and immunofluorescence (Figure 1C).

**Figure 1.**
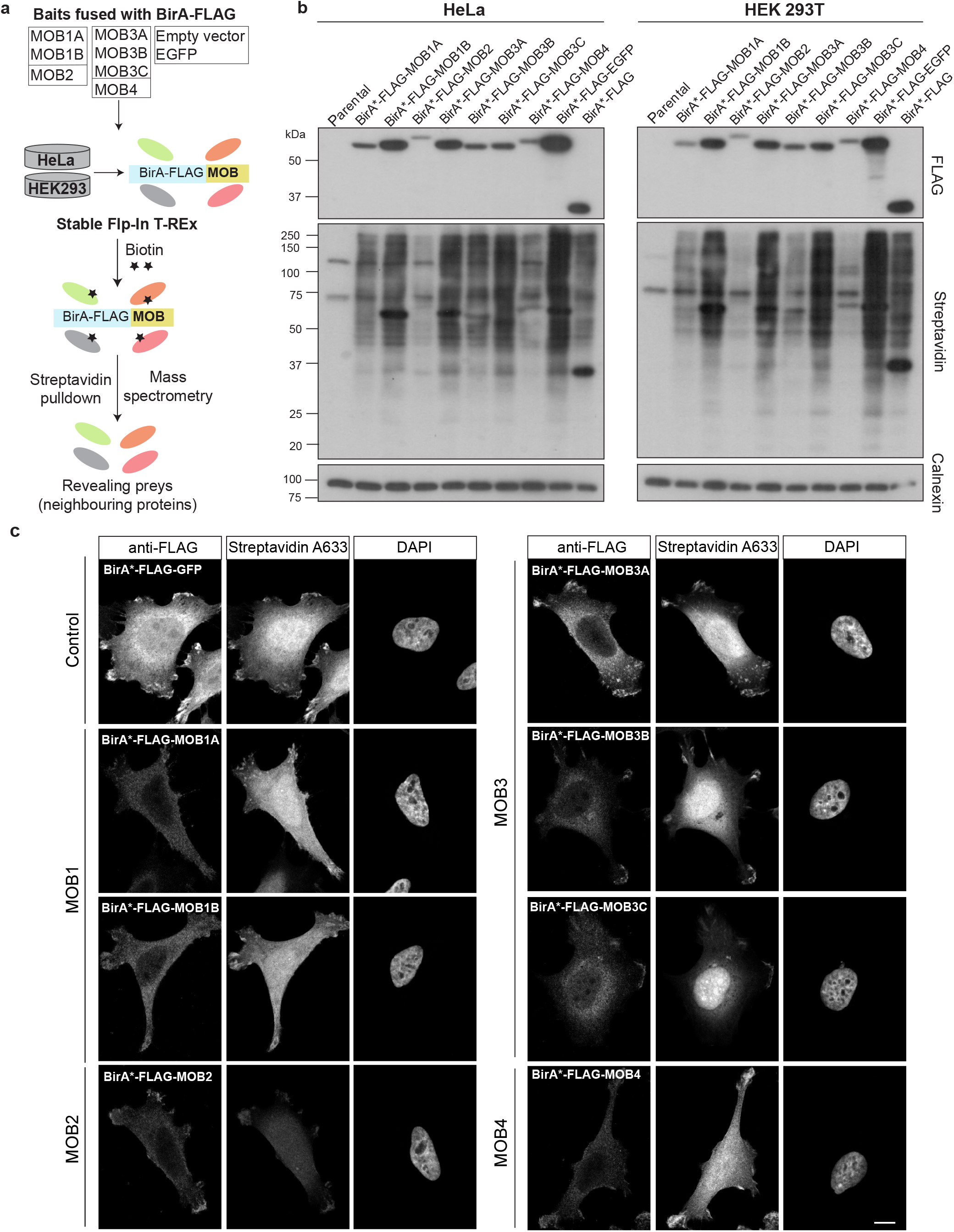
BioID proximity labelling screens for the MOB proteins. (a) Schematic outline of the BioID screens’ pipeline. (b) Western blot analysis of the parental and BioID bait-expressing Flp-In T-REx HeLa and HEK 293T cells. (c) Confocal microscopy images of the bait’s expression (anti-FLAG) and biotinylation (Streptavidin) patterns for the seven MOB proteins and a control condition (cells expressing BirA-Flag-EGFP) in HeLa cells. In (b-c), cells were treated with tetracycline (to induce expression) and biotin (to induce biotinylation) for 24 hours.

The BioID screens revealed 226 proximity interactors for the MOB family, 54 of which were shared between HeLa and HEK293 cells (Figure 2A, Table S2). Such overlap between the two cell lines (54/226 interactors; 24% of the dataset size) is in line with previously reported datasets for different proteins (20). Among the revealed interactions, 62 (27%) were previously reported in BioGrid (Figure 2B). These previously mapped interactions were mainly for MOB1A/B (48 interactions) and MOB4 (12 interactions) with none for MOB3A/B/C. MOB1s are an integral part of the Hippo pathway where the kinases MST1/2 phosphorylate MOB1 that binds to the kinase LATS allowing its autophosphorylation on the activation segment. MST1/2 also phosphorylate the hydrophobic motif of LATS for full activation. LATS phosphorylates and inhibits the activity of YAP1, the Hippo pathway effector (26). Indeed, we found that in both HEK293 and HeLa cells, all the pathway’s core components (LATS1/2 and STK3/4) and phosphatase (PP6 holoenzyme) were retrieved (16) (Figure 2C). Furthermore, the previously reported cytoskeleton-associated DOCK6-8 and LRCH1-3 proteins were also recalled (16). Similarly, the MOB2 *bona fide* interactors NDR kinases, STK38 and STK38L, were captured among top hits for MOB2 in our datasets (3,27) (Figure 2C). These findings gave confidence that our screens could reliably identify new protein interaction partners of MOB3 proteins, our primary focus.

**Figure 2.**
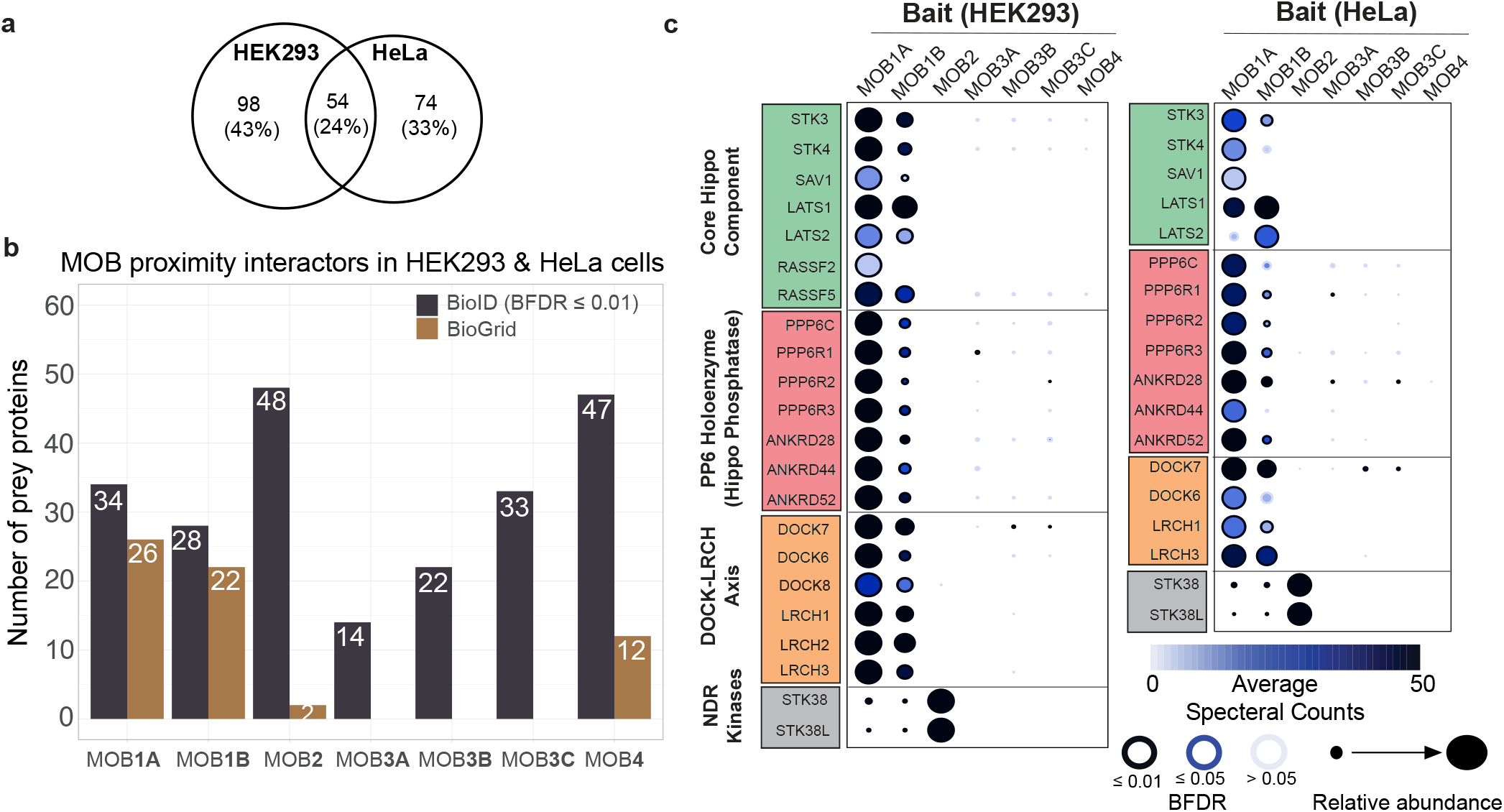
BioID recalls the *bona fide* MOB proteins interactors and extends their proximity network. (a) Venn diagrams showing the number of MOB preys identified in the two cell lines used in BioID. (b) Histogram demonstrating an enumeration of interactors identified for every MOB protein by either BioGrid or our BioID screens (BFDR ≤0.01). (c) Dot plots highlighting the *bona fide* interactors of MOB1A/B and MOB2 in HeLa and HEK293 cells.

### Global overview of the MOBs’ interactomes

Towards a holistic and individual MOB protein-based analysis, we applied a method that assigns a specificity score (MOB Specificity Score, MoSS; see Methods) for each prey towards a MOB bait after combining the datasets of the two cell lines together. This approach reassuringly pinpointed the Hippo pathway components and the Striatin (STRIPAK complex) as top specific interactors for MOB1A/B and MOB4, respectively, as outlined above and established in previous literature (11,16) (Figure 3A). Exploiting the MoSS metric, we generated an UpSet plot to globally compare the interactome of all the MOBs combined (Figure 3B). The highest number of shared proteins between MOBs was 15 proximal proteins between MOB1A and MOB1B. We then turned our focus to the MOB3 subfamily and first assessed whether they share neighboring proteins with other MOB subfamilies. The MOB3 subfamily shared only two proximal proteins (MAP4K4 and PTPN14) with the MOB1A/B subfamily (Figure 3C). Interestingly, both proteins are non-canonical Hippo pathway regulators, beyond the core components outlined above. A kinome-focused screen for MST1/2-independent kinases of LATS1/2 revealed six candidate kinases, including MAP4K4 that was further demonstrated to phosphorylate the hydrophobic motif of LATS and consequently inactivate YAP (28). Meanwhile, PTPN14, a nonreceptor protein tyrosine phosphatase, associates with and inhibits YAP’s activity (16,29–31).

**Figure 3.**
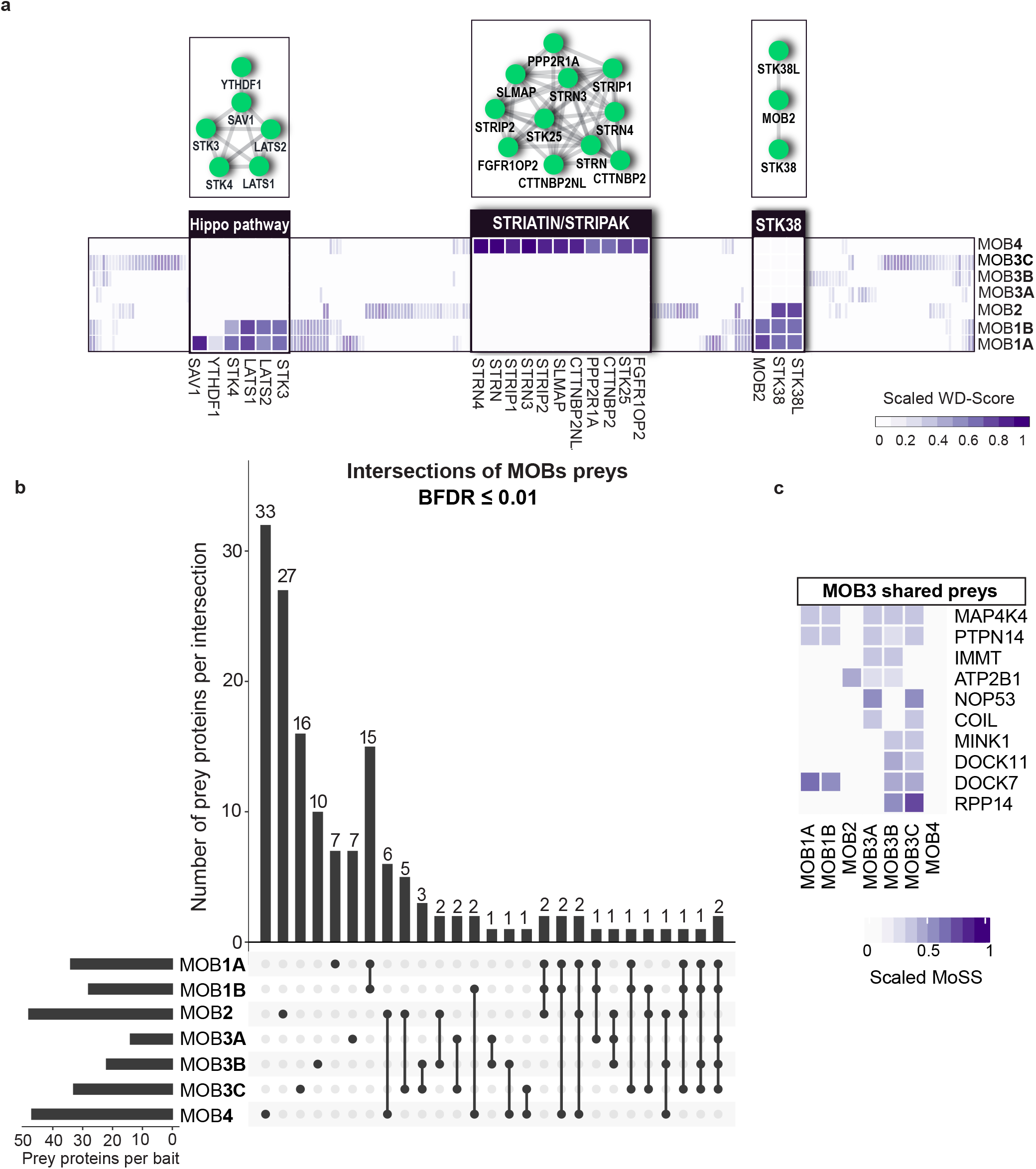
Overview of the interactome of the MOB subfamilies. (a) Heatmap highlighting the relative MoSS score for the *bona fide* interactors of MOB1A/B, MOB2, and MOB4. (b) An UpSet plot enumerating intersections of non-redundant proximal interactions identified for different MOB baits in either HeLa or HEK293 cells by BioID. (c) Heatmap highlighting the preys shared between MOB3A/B/C and other MOB subfamilies in addition to their individual common preys.

We then assessed whether MOB3A, MOB3B, and MOB3C share additional neighboring proteins. MOB3A shared IMMT, a subunit of the mitochondrial contact site and cristae organizing system (MICOS) complex (32), with MOB3B. MOB3A and MOB3B further shared the calcium transporter/pump ATP2B1(33) with MOB2. However, these two shared candidate interactors were captured at low abundance (Figure 3C, Table S2). MOB3A further uniquely shared two nuclear proteins, NOP53 and COIL, with MOB3C (Figure 3C). NOP53 is involved in ribosomal biogenesis and regulating the P53 pathway (34,35) while COIL is an integral/scaffold protein of the Cajal bodies and consequently mediates the assembly of small nuclear ribonucleoproteins (snRNPs) and their related essential functions such as mRNA splicing (36,37).

MOB3B and MOB3C shared four additional proteins, three just between themselves (MINK1, DOCK11, RPP14) and one (DOCK7) with MOB1A/B. MINK1 is implicated in different signaling networks such as mTORC2 and STRIPAK and was suggested as another LATS kinase (28,38,39). DOCK7/11 are guanine nucleotide exchange factors (GEFs) that control RAC1 and CDC42 to regulate cytoskeletal dynamics (40). Importantly, the connection between DOCK7 to MOB1A/B was described before (16). RPP14 is a protein subunit of the RNase P complex (see below). Taken together, these data demonstrate uniqueness of the interactome of the MOB3 subfamily.

### BioID reveals novel proximity interactions of MOB proteins

Towards defining candidate novel interactors of the MOB proteins and exploring uncharacterized members, we undertook two approaches: (i) defining preys that were not recalled from the BioGrid database (Table S2), and (ii) collective functional analysis of the proximity interactome of each MOB to explore whether there is an enrichment for specific protein complexes, cellular components, signaling pathways, and/or a specific biological process (Table S3). Expectedly, given the high number of previously defined interactors of MOB1A/B (Figure 2B), we did not define novel potential interactors that passed the stringent selection thresholds with respect to spectral counts and statistical significance (Figure 4A).

**Figure 4.**
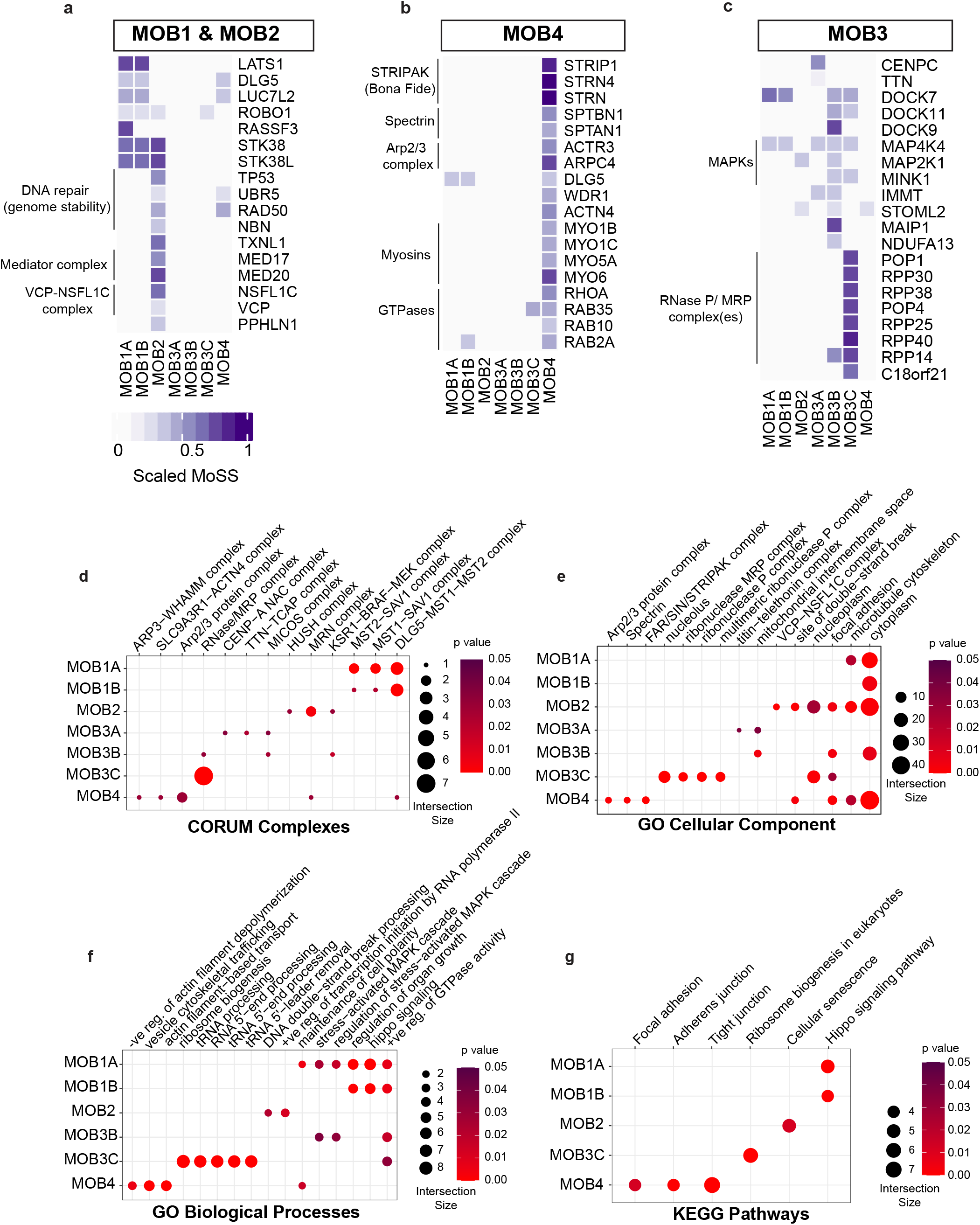
BioID reveals novel interactors for the MOB proteins. (a-c) Heatmaps highlighting proteins in the proximity of MOB1A/B, MOB2 (a), MOB4 (b), and MOB3A/B/C (c). (d-g) Dot plots displaying selected terms from overrepresented CORUM complexes (d), Gene Ontology (GO) Cellular Components (e), GO Biological Processes (f), and KEGG pathways (g).

MOB2 has recently been connected to DNA damage response and repair based on its interaction with RAD50 (41,42), which belongs to the MRE11A-RAD50-NBN (MRN) complex (43). This notion is reinforced in our dataset and extended further by capturing RAD50 and NBN in addition to TP53 and UBR5 in the vicinity of MOB2 (Figure 4A and 4D). Also, as part of the DNA damage response, we identified TXNL1, that regulates levels of the DNA repair protein XRCC1 (44–46), as a MOB2-specific prey (Figure 4A). The collective functional analysis of MOB2 interactome recalled components of the HUSH and VCP-NSFL1C complexes (Figure 4D-E). Periphilin (PPHLN1) forms the HUSH complex along with TASOR and MPP8 to mediate epigenetic silencing (47,48). VCP and its cofactor NSFL1C are required for membrane fusion and consequently the Golgi reassembly/regeneration (49–51). We identified PPHLN1, VCP, and NSFL1C as MOB2-specific prey proteins (Figure 4A). These observations collectively confirm the diverse nature of the defined MOB2 proximity interactome.

MOB4 proximity interactome was enriched for protein complexes, cellular components and functions closely tied to the cytoskeleton and its regulation (Figure 4D-G). For instance, we identified SPTAN1 and SPTBN1 (Figure 4B and 4E), two subunits of the heterodimeric Spectrin protein that organize(s) the cytoskeleton among other functions (52,53). We also defined members of the actin-based motor myosin protein superfamily (MYO1B/C, MYO5A, MYO6) (Figure 4B) that was further reflected in highlighting actin-based transport and trafficking as overrepresented biological processes in MOB4 interactome (Figure 4F). While these highlighted preys were not recalled on BioGrid as MOB4-interactors (Table S2), some of them demonstrated a specificity pattern towards MOB4 similarly to its *bona fide* interactor(s), STRIP1 and STRNs (Figure 4B).

Functional analyses did not pinpoint remarkable enrichments in the MOB3A-specific interactome (Table S3). In contrast, MOB3B interactome recalled the MAPK signaling pathway as evident by capturing MAP4K4 (Figure 4C and 4F). Remarkably, a strong association between MOB3C and the RNase P complex or the closely related mitochondrial RNase P (MRP) complex was suggested in all the performed analyses. The RNase P and MRP are ribonucleoprotein complexes, each of which is composed of a catalytic RNA and 10 protein subunits (8 of them are shared between the two complexes) to process pre-tRNA and pre-rRNA, respectively (54,55).

We found that 7 of the shared protein subunits between these complexes (POP1, POP4, RPP14, RPP25, RPP30, RPP38, RPP40) are exclusively among the top candidates in proximity to MOB3C (Figure 4C). Moreover, the complex(es) and its related functions such as ribosomal biogenesis and tRNA processing were recalled with high representation and statistical significance in all the functional analyses (Figure 4D-G, Table S3). Independent analyses of the two screened cell lines suggested that these candidates are the top proximity hits for MOB3C (Figure S1A-B).

In summary, our BioID data robustly revealed the diversity of the MOB proteins in terms of spatial organization and functions of their potential interactors. As a proof of principle for the described proximity network, we chose to further validate the MOB3C-RNase P interaction experimentally given that MOB3C is currently of unknown function and that no previous connection between MOB proteins and the RNase P/MRP has been reported.

### MOB3C physically interacts with catalytically active RNase P complex

We opted to validate the predicted interaction between MOB3C and the protein subunits of RNase P by performing co-immunoprecipitation experiments with individual subunits of the RNase P. Upon immunoprecipitating 3x-FLAG-MOB3C, we were able to detect YFP-POP1 but not YFP-RPP30 (Figure 5A), the two subunits with the highest abundance in the MOB3C interactome (Figure S1A-B). Furthermore, we failed to recover MOB3C in POP1 or RPP30 immuno-precipitates (Figure 5A). To count for the possibility that the potential interaction between MOB3C and these complexes happens with the assembled holoenzyme (i.e., the complete RNP complex), we performed an affinity purification-mass spectrometry (AP-MS) approach that included a chemical crosslinking step to validate the interaction. We generated tetracycline-inducible HeLa Flp-In T-REx cells expressing either 3xFLAG-MOB3C or 3xFLAG (Figure 5B) and induced the expression of MOB3C for 24 hours followed by a 2-hour treatment of dithiobis (succinimidyl propionate) (DSP) to crosslink the interactors before immunoprecipitation and MS analysis (Figure 5B). The different replicates from all the conditions showed robust tightness as suggested by high Spearman correlation score (Figure S2), giving confidence in the generated dataset. Towards our central question of whether MOB3C interacts with the RNase P protein subunits, we compared the BioID and AP-MS (with cross-linking) datasets and found only 10 shared proteins. Among those 10 were the 7 RNase P/MRP protein subunits, that notably were not retrieved in the AP-MS condition without crosslinking (Figure 5C and Table S4). Furthermore, the RNase P complex and its tRNA processing function were identified among the top represented CORUM complexes and Gene Ontology Biological Processes in the cross-linked AP-MS dataset, respectively (Figure S3A-B and Table S5). Overall, these results confirmed that MOB3C interacts with the RNase P RNP complex.

**Figure 5.**
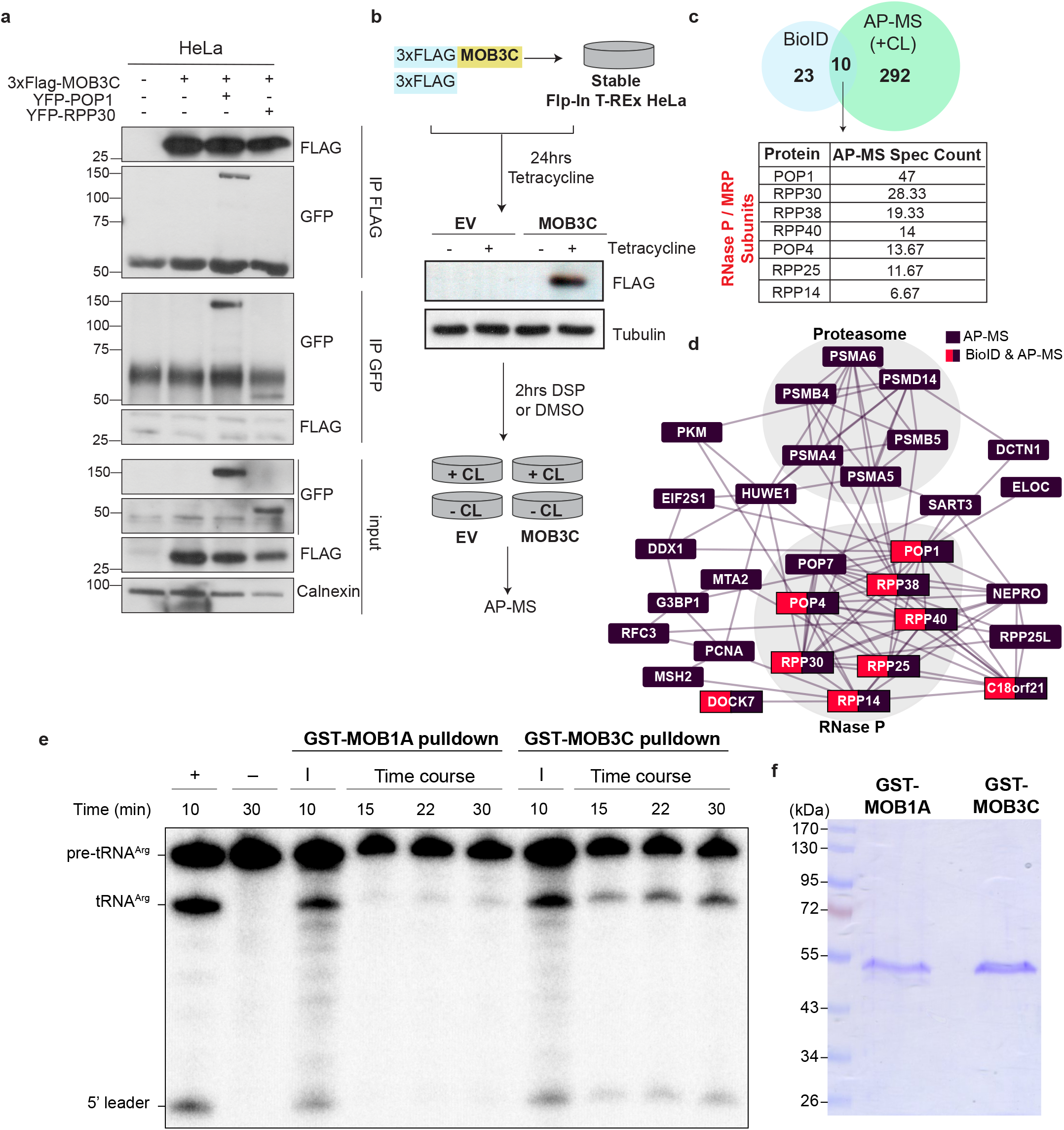
MOB3C interacts with catalytically active RNase P complex. (a) Co-immunoprecipitation assay for YFP-tagged POP1 and RPP30 with FLAG-tagged MOB3C in HeLa cells. (b) Schematic outline for the affinity purification-mass spectrometry (AP-MS) with DSP cross-linking. A western blot analysis validating the used cell lines in AP-MS is shown. EV, empty vector; DSP, dithiobis (succinimidyl propionate); CL, crosslinking. (c) Venn diagram showing the commonly defined proteins in the BioID and cross-linked AP-MS datasets. The table displays the average spectral counts of each RNase P subunit defined in the AP-MS dataset. (d) A proximity network of the RNase P protein subunits defined in both the MOB3C BioID and AP-MS datasets in reference to BioGrid (see Methods). (e) Pre-tRNA cleavage assay of the pulldown experiments using GST-MOB1A or GST-MOB3C. Time-course reactions showing RNase P activity when GST-MOB3C, but not GST-MOB1A, was used. +: a positive control of pre-tRNA^Arg^ cleaved by recombinant *E. coli* RNase P; –: a negative control of pre-tRNA^Arg^ without enzyme; I: Input. (f) SDS-PAGE analysis confirms the presence of GST-MOB3C and GST-MOB1A post-pulldown.

The RNase P and its subunits might interact with different proteins or complexes to mediate different functions beyond the canonical tRNA processing role (56,57). This notion is only starting to be unravelled by building the RNase P proximity network; therefore, we asked whether the MOB3C BioID and crosslinked AP-MS datasets can intersect with such a network. Interestingly, our assembled network recalled previously identified interactors of the RNase P complex or at least some of its subunits such as NEPRO (58), PCNA, RFC, MSH2, and different subunits of the proteasome (56,59) (Figure 5D). Notably, within this network was C18orf21, that was identified as one of the 10 common prey proteins between the MOB3C AP-MS and BioID.

To assess whether MOB3C interacts with a functional RNase P complex, we pulled down RNase P using GST-MOB3C or GST-MOB1A (negative control) as baits and assayed for precursor tRNA cleavage activity. We observed that significant pre-tRNA^Arg^ (n-Tr21) cleavage activity was pulled down by MOB3C compared to the MOB1A negative control (Figure 5E). We also confirmed that both GST-MOB3C and GST-MOB1A were pulled down with similar efficiency by glutathione beads (Figure 5F). Collectively, these results strongly suggest that MOB3C interacts with catalytically active RNase P holoenzyme.

## DISCUSSION

The MOB proteins, at least the well-studied family members, mediate their functions through engaging in protein-protein interactions and assembling complexes. Here, we systematically uncovered the proximity network of the MOB family and found a minimal overlap between different subfamilies (e.g., MOB1 versus MOB3). Importantly, the interactomes of the 3 MOB3 proteins are different, arguing against considering them as a subfamily with a unified function (8,15). MOB3A and MOB3C are upregulated in gliomas and combined depletion of all three human MOB3 proteins halted the proliferation of a glioblastoma cell line *in vitro* and *in vivo* (15). The study concluded that the human MOB3s mediate a pro-tumorigenic effect. However, the authors did not assess the effect of individual depletions of MOB3A/B/C, but instead depleted simultaneously all 3 MOB3s, assuming they offer functional redundancy. Our unbiased proteomic screens argue against such a notion.

In addition to comparing the subfamilies to each other, our screens identified novel interactors for different MOBs that potentially guide the search for their molecular functions. Importantly, BioID showed superiority in revealing such interactions when compared to traditional methods, such as the yeast two-hybrid assays used to reveal MOB2 interactors or AP-MS strategy used with human MOBs (17,41). Some of the identified interactions extend previous findings (e.g., connecting MOB2 to DNA repair proteins, and MOB4 to cytoskeletal remodelers), while others spawn new directions (e.g., connecting different MOBs to MAPKs that might potentially converge on the Hippo pathway). The latter feeds into the unresolved relationship between MOB3 proteins and this pathway. In agreement with our BioID screens, previous co-immunopreciptation assays or AP-MS screens suggested that MOB3 proteins do not interact with or activate LATS or STK38 (17,27). Meanwhile, an interaction between MOB3 proteins and the Hippo pathway core kinase, MST1, was reported only under specific experimental conditions (8,15), which our screens did not recall. Here, we report the existence of the non-canonical Hippo regulators MAP4K4/6 and PTPN in the vicinity of MOB3 proteins. However, such notions would require thorough experimental validations in future studies, since proteins defined in BioID screens are not necessarily direct interactors with the bait protein.

The biological function of MOB3C remains elusive. Recently, it was suggested to facilitate bypassing oncogene-induced senescence (8). However, these observations were obtained by studying a myristoylated form of MOB3C, which might only account for a small fraction of the native MOB3C. Here, through unbiased proteomics and subsequent GST-pulldowns, we reveal an unexpected connection between MOB3C and RNase P/MRP, two evolutionarily conserved and essential ribonucleoprotein complex(es). This observation rationalizes multiple questions. First, does the MOB3C-RNase P interaction require other intermediary proteins? Second, because the seven RPPs identified in the vicinity of MOB3C are shared between the RNase P and RNase MRP (60), does MOB3C interact preferentially with both RNPs? In both cases, the reason and consequences behind this interaction should be addressed by future studies. One possibility is MOB3C regulating the complex(es)’ catalytic activity or substrate specificity. This would go in line with the view that the H1 RNA of the RNase P accumulated additional protein subunits through evolution to tune its affinity towards different substrates (24,61,62). Alternatively, MOB3C might act as an adaptor to help assemble the holoenzyme and hence raises the question of which subunit(s) it binds. The specificity of the RNase P complex for MOB3C was clear in our BioID interactomes, a convincing demonstration that individual MOB3 proteins have evolved distinct functions. While the proteins share 70-80% identity at the amino acid level and the AlphaFold predicted structures are indistinguishable (average backbone r.m.s.d. of 0.25Å), MOB3C has acquired several additional charged residues, particularly arginine, across a broad region of the protein that includes an unstructured loop that could complex with a ribonucleoprotein such as RNase P. The 12 arginine residues on this face of MOB3C are significantly more than those found in MOB3A (6) or MOB3B (4), a potential rationalization for their differential binding to the RNase P complex. Future work is needed to test which complex is indeed interacting with MOB3C and the functional consequences.

In summary, our study not only provides a resource for exploring new interactors and consequently functions of the seven human MOB proteins, but also puts forward an interesting connection between the MOB family and RNA processing.

## METHODOLOGY

### Cell lines and culture

HEK293 Flp-In T-REx cells were purchased from ThermoFisher Scientific, while HeLa Flp-In T-Rex were a gift from S. Taylor (University of Manchester, UK). All cell lines were cultured in Dulbecco’s modified Eagle’s medium (DMEM; Wisent #319-005-CL) supplemented with 10% fetal bovine serum (FBS) and 1 % penicillin/streptomycin (P/S) antibiotics (Wisent).

### Generating stable cell lines and BioID pipeline and analysis

To generate Gateway-compatible sequences, the 7 human MOBs complementary DNA sequences (including the stop codons) flanked by the attB sequences were synthesized (Twist Bioscience). These cDNAs were consequently recombined into the pDONR-221 vector and then shuttled into the pcDNA5-FRT backbone vector expressing an abortive mutant of BirA (BirA*) tagged with FLAG using the Gateway recombination cloning system (16,63). Generating the stable Flp-In T-REx expressing BirA*-FLAG-MOBs or BirA*-FLAG or BirA*-FLAG-EGFP was done according to the manufacturer protocol and as described here (63). For validating the construction of the cell lines, cells were seeded into 6 well-plates (Corning) and incubated overnight. The following day, cells were induced with tetracycline (1 µg/ml) and/or treated with biotin (50 µM) for 24 hours. Cells were then lysed with ice-cold Radio ImmunoPrecipitation Assay (RIPA) buffer (1% Nonidet P-40, 0.5% deoxycholic acid, 0.1% SDS, 150 mM NaCl, 5 mM EDTA, 50 mM Tris pH 7.5) supplied with NaF (5 mM), Na3VO4 (1mM) and 1x complete protease inhibitor (Roche). Western blotting was done using the following concentrations of the antibodies: monoclonal anti-FLAG M2-peroxidase (HRP) (1:10000; Sigma #A8592) and streptavidin-HRP (1:25,000; BD Biosciences #554066) in Tris-buffered saline (TBS) solution (1% bovine serum albumin and 0.1% Tween-20).

For BioID, cells were seeded in 15 cm plates (Sarstedt) to reach 70-80% confluence. Subsequently, simultaneous induction with tetracycline (1 µgml^−1^) and treatment with biotin (50 µM) was done 24 hours prior to processing the cells as we previously detailed (20,63) with no modifications.

Mass spectrometry data were analyzed as we previously described (20). Briefly, the raw data were analyzed with Prohits (64). The MOBs’ two datasets (HEK293 and HeLa) were compared to their respective controls (BirA-FLAG and EGFP-BirA-FLAG). To define interaction statistics, we used SAINTexpress (v3.6.1) (65) through ProHits (64) by using the following parameters: iProphet protein probability ≥ 0.9 and unique peptides ≥ 1. Each of the proteomics datasets (HEK293 and HeLa) was compared individually versus their negative controls (EGFP and empty vector). SAINT analyses were performed using the following settings: nControl:3, nCompressBaits:3 (no baits compression). Preys’ baits specificities (WD-score) were calculated for each cell lines separately by using the CompPass algorithm (66) after we uploaded the SAINT results to the Prohits Prey Specificity online tool (https://prohits-viz.lunenfeld.ca/Specificity/). Interactions having a BFDR ≤ 0.01 were considered as statistically significant. Next, we estimated the MOB specificity score (MoSS) index for each prey by summing both of their cell lines WD-scores. The data were log2-transformed and rescaled on a range of 0.1 to 1.0 in R (www.r-project.org) with the Scales package. After we assigned a value of zero to unidentified interactions, we created a heatmap with the pheatmap package in R. An UpSet plot was created with the UpSetR package in R by enumerating non-redundant interactions identified by BioID from both HEK293 and HeLa cell lines. Enumerations of interactions were calculated on statistically significant preys (BFDR ≤ 0.01) identified in both cell lines for all seven MOB baits. We also enumerated interaction recalls by identifying within our statistically significant preys known MOBs interactions from the human BioGrid database (version 4.4.208). A histogram of these enumerations was created in R by using the ggplot2 package. The R package gprofiler2 was used to analyze the overrepresented GO terms, KEGG pathways and CORUM complexes (67) by statistically correcting p-values with the fdr correction method. We selected statistically significant terms having adjusted p-values < 0.05 and presented the results in dotplots using the ggplot2 R package. The BioID and AP-MS results were imported in Cytoscape (v.3.9.1) (cytoscape.org) to build graphical representations of protein–protein interaction networks. We then performed a network augmentation by extracting prey–prey interactions from the human BioGRID database (v.4.2.192) (68) and from Cytoscape’s PSICQUIC Web Service client (May 2021 release) through the IntAct and UniProt databases.

#### Dot plot analyses

Dot plots were produced through the ProHits-Viz online tool (https://prohits-viz.org/) with the generated SAINT output files, using a BFDR ≤ 0.01 as a statistical cut-off.

### Affinity purification-mass spectrometry

MOB3C cDNA was shuttled into the pDEST 5’ Triple FLAG pcDNA5-FRT to generate an N-terminally FLAG-tagged MOB3C. Both this vector and the original pDEST5’ Triple FLAG pcDNA5-FRT vector were independently used to generate stable Flp-In T-REx HeLa cells expressing either FLAG-MOB3C or FLAG (empty vector), respectively (63). These two cell lines were seeded in 6 well plates overnight then induced with tetracycline (1 µgml^−1^) for 24 hours and then processed as described earlier to validate the expression of MOB3C.

For the AP-MS experiments, the generated Flp-In T-REx HeLa cells were seeded in 15 cm plates overnight and induced with tetracycline (1 µgml^−1^) for 24 hours before processing for the AP-MS. For the crosslinking, DSP (ThermoFisher Scientific #22585) was dissolved in DMSO for a 100 mM stock solution and a warm 1% DSP solution in calcium-and magnesium-free Dulbecco’s phosphate-buffered saline (DPBS; ThermoFisher Scientific #21600010) was added on the cells for 2 hours on an ice-water bath for crosslinking (69). Subsequently, cells were incubated for 15 minutes on ice with a DSP quenching solution (20 mM Tris pH 7.5 in DPBS). The AP-MS experiments were done as previously reported (16) with the following modifications. Briefly, cells were collected and lysed in 0.1% NP40 lysis buffer (100 mM KCl, 50 mM Hepes-KOH pH 8.0, 2 mM EDTA, and 10% glycerol) supplemented with 1x protease inhibitor cocktail, 1mM DTT and 1 mM PMSF. Affinity purification was carried out using 20 μl of anti-FLAG M2 magnetic beads (Sigma; #M8823-1ML) per condition. The on-bead proteins were first diluted in 2M Urea/50mM ammonium bicarbonate and on-bead trypsin digestion was performed overnight at 37°C. The samples were then reduced with 13mM dithiothreitol at 37°C, cooled for 10 minutes and alkylated with 23 mM iodoacetamide at room temperature for 20 minutes in the dark. Trifluoroacetic acid was used to acidify supernatants. MCX cartridges (Waters Oasis MCX 96-well Elution Plate) were then used to clean the supernatants from residual detergents and reagents according to the manufacturer’s instructions. After elution in 10% ammonium hydroxide /90% methanol (v/v), samples were dried in a Speed-vac, reconstituted under agitation for 15 min in 22 µL of 2%ACN-1%FA (formic acid), and loaded onto a 75 μm i.d. × 150 mm Self-Pack C18 column installed in the Easy-nLC II system (Proxeon Biosystems). The solvents used for chromatography were 0.2% formic acid in water (solvent A) and 0.2% formic acid in acetonitrile (solvent B). Peptides were eluted with a two-slope gradient at a flowrate of 250 nL/min. Solvent B first increased from 2 to 34% in 80 min and then from 34 to 80% B in 12 min. The HPLC system was coupled to an Orbitrap Fusion mass spectrometer (Thermo Scientific) through a Nanospray Flex Ion Source. Nanospray voltage was set to 1.3-1.7 kV, meanwhile the S-lens voltage was set to 60 V. Capillary temperature of 225 °C was set. Full scan MS survey spectra (m/z 360-1560) in profile mode were acquired in the Orbitrap with a 120,000 resolution and a 3e5 target value. The 25 most intense peptide ions were fragmented in the HCD collision cell and analyzed in the linear ion trap with a target value at 1e4 and a normalized collision energy at 29. Target ions selected for MS/MS fragmentation were dynamically excluded for 20 sec after two MS2 events.

### Immunostaining and confocal microscopy

HeLa Flp-In T-REx cells were plated on coverslips and simultaneous induction with tetracycline (1 µgml^−1^) and treatment with biotin (50 µM) was done 24 hours prior to cell fixation with 3.7% formaldehyde in CSK buffer [100 mM NaCl, 300 mM sucrose, 3LmM MgCl_2_, 10 mM PIPES, pHL6.8]. Then, cells were permeabilized with 0.2% Triton X-100 in PBS for 5 min before incubation with rabbit polyclonal FLAG antibody (Cell signaling technology #2368) 1:1000 dilution in the wash buffer [1%LBSA, 0.1% Triton-X100 in TBS], rinsed 5 times with wash buffer and incubated for 30 min with Alexa Fluor 488 conjugated chicken anti-rabbit (Life Technologies #A-21441) 1:500 dilution together with Hoechst 33342 (Invitrogen #H3570) 1:10 000 dilution and Alexa Fluor 633 conjugated streptavidin (Thermo Fisher Scientific #S21375) 1:500 dilution in wash buffer. Cells were washed ten times with the wash buffer and one time with water. Coverslips were mounted with Mowiol. Images were acquired with a Carl Zeiss LSM700 laser scanning confocal microscope (Carl Zeiss MicroImaging, Jena, Germany) equipped with a plan-apochromat 63x/1.4 numerical aperture objective and operated with ZenBlack 2009.

### Co-immunoprecipitation experiments

YFP-RPP30-C1 was a gift from Susan Janicki (Addgene plasmid #134547). YFP-POP1 expression vector was generated by amplifying the POP1 cDNA from pCMVh-POP1-3xFLAG (gift from Martinn Dorf; Addgene #53968) with flanking XhoI and SalI restriction sites and ligating it into the multicloning site of YFP-C1 backbone vector by swabbing out the YFP-RPP30. Primers used: XhoI-POP1 (Forward:taagcaCTCGAGaaatgtcaaatgcaaaagaaag) and POP1-SalI (Reverse: tgcttagtcgactcacacctcaatagcaatcctcg). Flp-In T-REx HeLa expressing FLAG-MOB3C were transfected with 10 ug of either YFP-RPP30 or YFP-POP1 using Lipofectamine 2000 (Invitrogen) according to the manufacturer’s instructions. 24 hours after transfection, the expression of MOB3C was induced with tetracycline (1 µgml^−1^) for 24 hours. 48 hours after transfections and 24 hours after tetracycline induction, cells were lysed with CHAPS lysis buffer (0.5% CHAPS, 1mM MgCl_2_, 1mM EGTA, 10mM Tris pH7.5, 10% glycerol, 5mM β-mercaptoethanol, and 5mM NaF) supplemented with 1x complete protease inhibitor and 1mM Na_3_VO_4_ as previously reported (70). For immunoprecipitation, 20µl/condition of either anti-FLAG M2 affinity gel (Sigma) or Protein G agarose beads (Genscript) were washed three times with 1 mL of immunoprecipitation buffer (70) (0.1% Triton X, 20 mM Tris pH 7.5, 10 mM MgCl2, 150 mM KCl, and 10% glycerol). Beads were then incubated for 2h at 4°C with 1 mg of protein lysate alone or with the addition of 1µl of GFP antibody (Life technologies) for the FLAG and GFP IP, respectively. Immunoprecipitates were then washed three times with the immunoprecipitation buffer and denatured in 6x sample buffer (350mM Tris-HCl pH6.8, 10% SDS, 30% glycerol, 0.1% Bromophenol blue, 10% β-mercaptoethanol). Lysates and immunoprecipitates were then processed for immunoblotting using the following concentrations: Flag M2-HRP 1:10000 (Sigma # A8592), GFP 1:2000 (Life technologies #A-11122) and Calnexin E-10 1:500 (Santa Cruz #sc-46669) in TBS solution.

### Purification of GST-MOB3C and GST-MOB1A

The plasmid pTH35-GST-MOB3C was used to transform Escherichia coli BL21 RIL Codon Plus (Agilent) cells. A single colony was used to inoculate 5 mL LB medium containing 100 μg/mL carbenicillin, 25 μg/mL chloramphenicol and 1% (w/v) glucose, and the culture was grown overnight at 37°C with shaking. This seed culture was used to inoculate 500 mL LB medium (supplemented with carbenicillin and chloramphenicol as above). The culture was incubated at 37°C with shaking until OD_600_ ∼ 0.6 before induction with 0.4 mM IPTG and growth continued at 15°C overnight. The cells were harvested by centrifugation (6000 g, 15 min, 4°C) and the pellet was re-suspended in 20 mL of buffer A (1x PBS (pH 7), 0.5 M NaCl, 1 mM PMSF, 1mM DTT). The resuspended cells were lysed by sonication and the cell lysate was cleared by centrifugation (23,000 g, 30 min, 4°C). The resulting supernatant was added to 800 μl of glutathione agarose (Thermo Scientific), which was equilibrated in lysis buffer. After nutating at 4°C for 1 h, the resin was pelleted at 700 g for 2 min and washed extensively with wash buffer (1x PBS (pH 7), 0.5 M NaCl, 1mM DTT), before eluting twice with 3 mL and then 2 mL of elution buffer (50 mM Tris-HCl (pH8), 150 mM NaCl, 1 mM DTT, 20 mM reduced glutathione). After confirming the presence of GST-MOB3C by SDS-PAGE, the eluents were pooled and diluted with buffer A0 [50 mM Tris-HCl (pH 7.5), 1 mM DTT] to reduce the NaCl in the sample to 25 mM. The sample was then loaded on a 1-mL Heparin HP Sepharose column (GE Healthcare), which was equilibrated with buffer A50 (50 mM Tris-HCl (pH 7.5), 50 mM NaCl, 1 mM DTT). The column was washed extensively with buffer A50 and the protein eluted with a linear 0–1 M NaCl gradient using buffer A1000 (50 mM Tris-HCl (pH 7.5), 2M NaCl, 1 mM DTT) and an AKTA FPLC purifier (GE Healthcare). After SDS-PAGE analysis, fractions containing GST-MOB3C were pooled and dialyzed overnight at 4°C against 1 L of buffer C (20 mM Tris-HCl (pH 8), 150 mM NaCl, 14.3 mM β-mercaptoethanol, 10% glycerol). The dialyzed sample was concentrated using a 5-kDa Amicon centrifugal filter (Milipore) with molecular weight cut off of 5kDa and the final protein concentration was determined by measuring Abs280 using a Nanodrop spectrophotometer and using an extinction coefficient of 75,750 M-1cm-1 (ExPASy ProtParam Tool) (71). The protein was flash frozen and stored at –80°C until use. A similar protocol was followed to obtain GST-MOB1A using plasmid pTH35-GST-MOB1A. The concentration of GST-MOB1A was determined using an extinction coefficient of 72,770 M-1cm-1 (ExPASy ProtParam Tool) (71).

### Pulldown of HeLa RNase P using GST-MOB3C and GST-MOB1A

HeLa cell pellets were harvested from confluent 15-cm plates and stored at -80°C until use. Cell pellets were gently resuspended in 750 μL of lysis buffer (20 mM Tris-HCl (pH 8), 100 mM NaCl, 5 mM MgCl2, 1% glycerol; 14.3 mM β-mercaptoethanol, 0.2 mM PMSF). One third of the resuspended cells was lysed by sonication, and the remaining two thirds were stored at -80°C for future use. The cell lysate was cleared by centrifugation (20,000 g, 20 min, 4°C). One-half of the supernatant was mixed with ∼120 pmol GST-MOB3C and the other half with ∼120 pmol GST-MOB1A. Both samples were incubated at 37°C (with nutation) for 15 min and then on ice for 30 min. Two μL of each sample were saved as the input (I). Each sample was then added to 80-μL glutathione agarose beads (Thermo Scientific), which had been pre-equilibrated in lysis buffer. All samples were nutated at 4°C for 90 min before the beads were harvested and washed twice with 500 μL of wash buffer (20 mM Tris-HCl (pH 8), 100 mM NaCl, 5 mM MgCl2, 14.3 mM β-mercaptoethanol, 0.005% (v/v) NP-40). One-third of the beads was used for RNase P activity assay with precursor n-Tr21 (pre-trRNA^Arg^) as the substrate (72). The beads were mixed with 10 μL reaction buffer (20 mM Tris-HCl (pH 8), 14.3 mM β-mercaptoethanol, 100 mM KCl, 1 mM MgCl2, 20 nM pre-n-Tr21, a trace amount of which was internally labeled with [α-32P]-GTP-labeled). The reaction was nutated at 37°C, and 3-μL aliquots were withdrawn at 15, 22, and 30 min and quenched with 10 μL loading dye (7 M urea, 20% (v/v) phenol, 0.2% (w/v) SDS, 5 mM EDTA, 0.05% (w/v) bromophenol blue and 0.05% (w/v) xylene cyanol). The reaction products were then separated on a denaturing PAGE gel (10% (w/v) polyacrylamide, 7 M urea), and visualized using a Typhoon Phosphorimager (GE Healthcare). One-fifth of the beads was analyzed by SDS-PAGE and Coomassie blue staining to confirm the presence of GST-MOB1A and GST-MOB3C in the precipitates. The pulldown experiment was independently repeated two times and the same trend was observed. Only one representative gel is shown.

## ACKNOWLEDGMENTS

We would like to thank the IRCM Proteomics facility staff for their technical help. This work was supported by operating grants from the Cancer Research Society (#25244 to J.-F.C), Henry M. Jackson Foundation for the Advancement of Military Medicine (subaward 5516 to V.G.), and the Behrman Research Fund (to V.G.). I.E.E. was a recipient of FRQS Doctoral and IRCM Foundation-TD scholarships, and H-D.P was supported by an OSU Comprehensive Cancer Center Pelotonia Pre-doctoral Fellowship. J.-F.C. holds the Canada Research Chair Tier 1 in Signalling in Cancer and Metastasis, and the Alain Fontaine Chair in Cancer Research and the Transat Chair in Breast Cancer from the IRCM Foundation.

## CRediT author statement

Islam Elkholi & Jean-François Côté: **Conceptualization**, **Supervision, Writing - Finalizing the manuscript**. Islam Elkholi, Jonathan Boulais, Marie-Pier Thibault, Hong-Duc Phan, Amélie Robert, Lien B. Lai: **Methodology** & **Investigation**. All authors: **Formal analysis** & **Writing - Review & Editing**. Islam Elkholi: **Writing - Original Draft**. Jean-François Côté: **Funding acquisition**.

## SUPPORTING INFORMATION

This article contains supporting information. The generated datasets (BioID and AP-MS) and their respective functional analyses are provided in the supplementary tables (S1-5).

---

**Figure S1.**
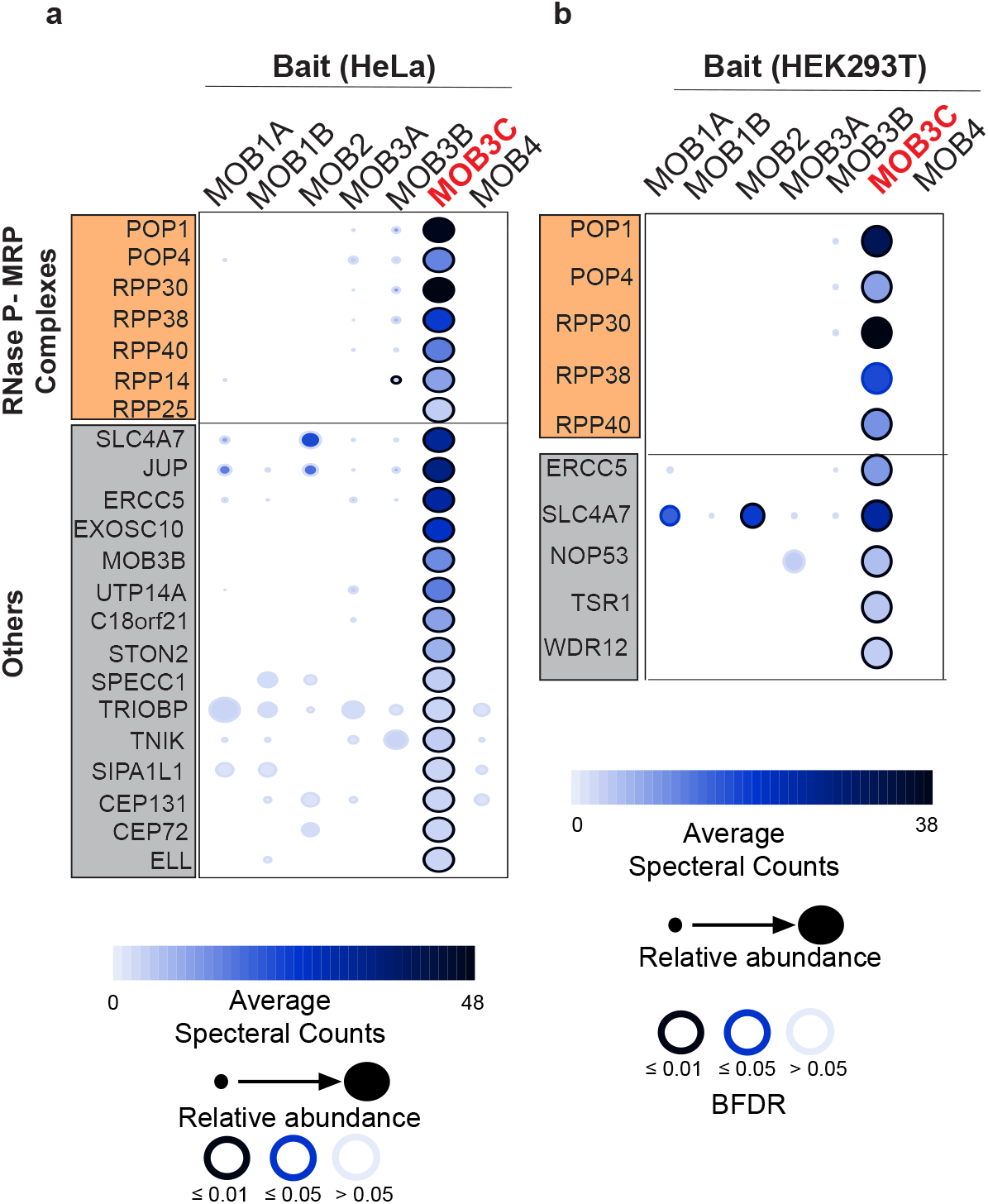
BioID reveals new interactors for MOB3C. (a-b) Dot plots showing the enrichment of the protein subunits of the RNase P/MRP complexes in the vicinity of MOB3C in HeLa (a) and HEK293 (b) cells.

**Figure S2.**
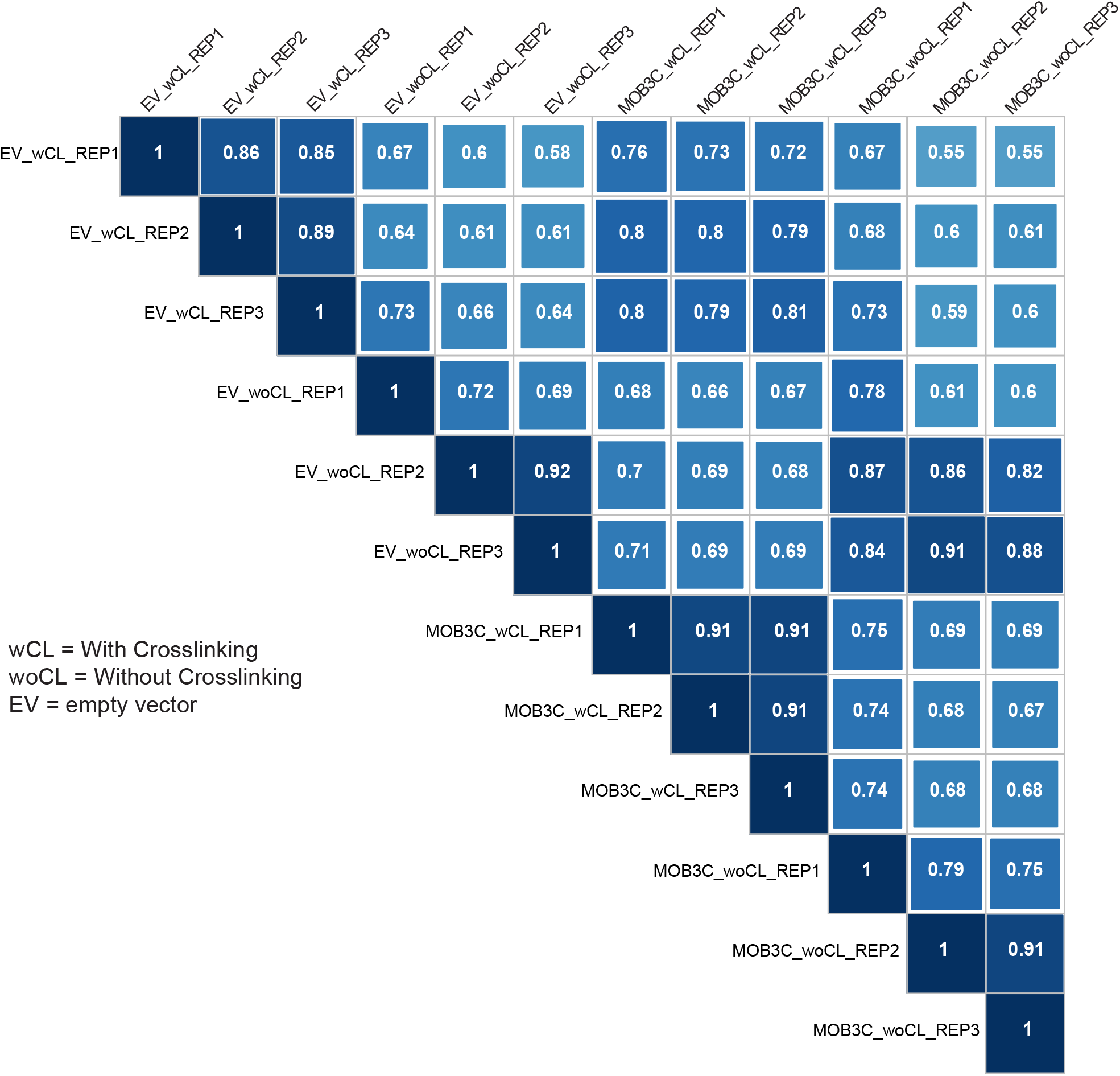
Quality control for the generated AP-MS datasets. The biological replicates of the DSP-crosslinked MOB3C condition showed high Spearman correlations (0.91) with each other confirming tightness between these replicates. REP, replicate.

**Figure S3.**
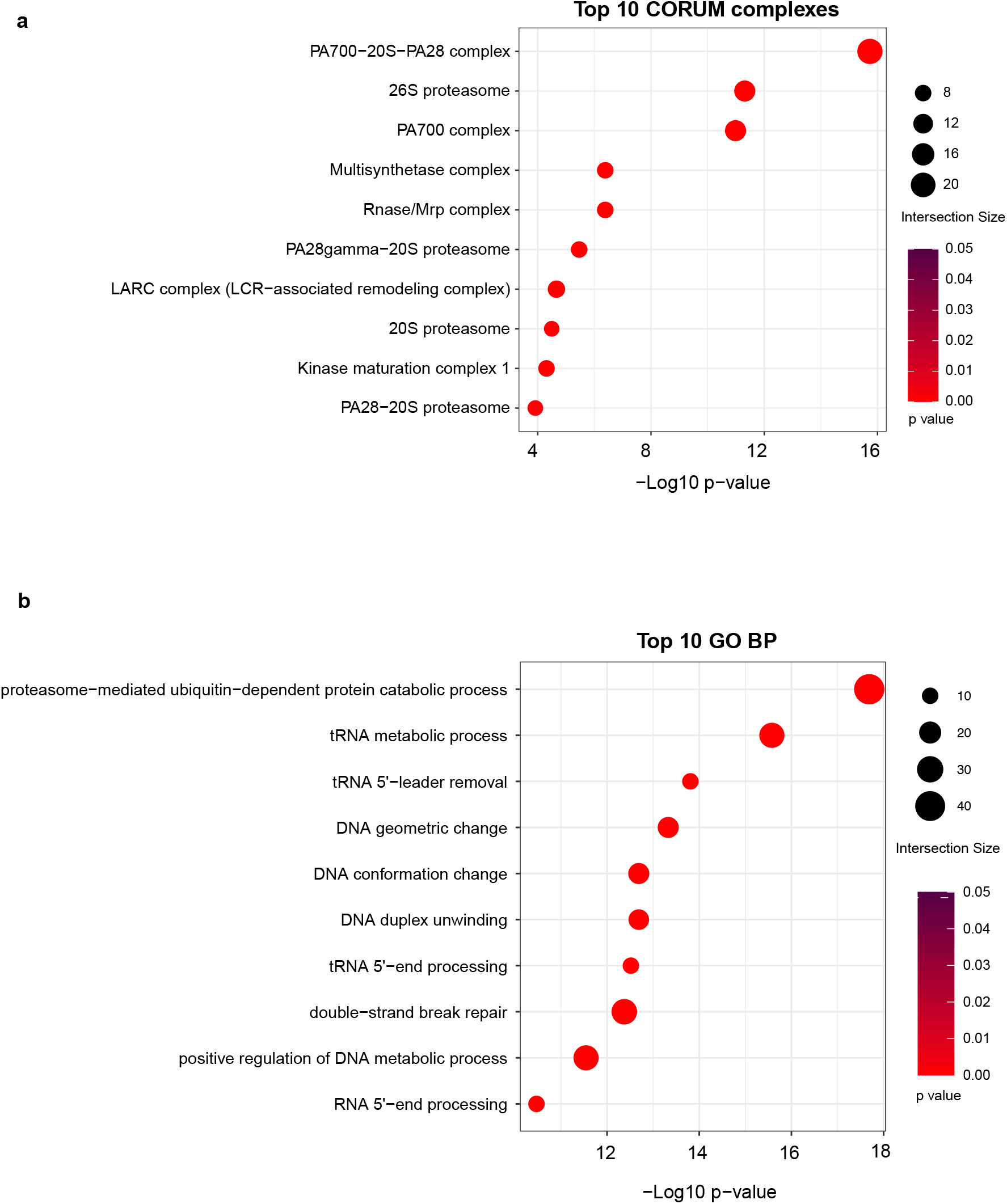
Functional analyses for MOB3C interacting proteins. The list of statistically significant interactors of MOB3C from the cross-linked AP-MS dataset was used to define the top 10 overrepresented CORUM complexes (by adj. *p*-values) (a), and Gene Ontology Biological Processes (BP) (b).

